# State-of-the-art EEG artifact removal evaluation

**DOI:** 10.1101/2021.10.23.465532

**Authors:** Zhenyu Jin

## Abstract

**Object:** Electroencephalography (EEG) signals suffer from a low signal-to-noise ratio and are very susceptible to muscular, ambient noise, and other artifacts. Many artifact removal algorithms have been proposed to address this problem. However, the evaluation of these algorithms is conventionally too indirect (e.g., black-box comparisons of brain-computer interface performance before and after removal) because it is unclear which part of the signal represents raw EEG and which is noise. This project objectively benchmarks popular artifact removal algorithms and evaluates the fundamental Independent Component Analysis (ICA) approach thanks to a unique dataset where EEG is recorded simultaneously with other physiological signals-facial electromyography (EMG), accelerometers, and gyroscope-while ten subjects perform several repetitions of common artifact-inflicting tasks (blinking, speaking, etc.).

**Approach:** I have compared the correlation between EEG signals and the artifact-representing channels before and after applying an artifact removal algorithm across the different artifact-inflicting tasks. The extent to which an artifact removal method can reduce this correlation objectively quantifies its effectiveness for the different artifacts. In the same direction, I have determined to what extent ICA successfully detects artefactual components in EEG by comparing the corresponding correlations for independent components that are labelled as artifacts with those labeled as EEG.

**Main result:** The FORCe was found to be the most effective and generic artifact removal method, cleaning almost 40% of artifacts. ICA is shown to be able to isolate almost 70% of artefactual components.

**Significance:** This work alleviates the problem of unreliable evaluation of EEG artifact removal frameworks and provides the first reliable benchmark for the most popular algorithms in this literature.

## Introduction

In recent years, with the developments in neuroscience, cognitive science, and cognitive psychology, electroencephalography (EEG), functional near-infrared spectroscopy (fNIRS), magnetoencephalography (EMG), and other neuroimaging tools have gained much attention. As a tool for analyzing brain activity, EEG signals can record from multiple electrodes on the scalp [1] to reflect the functional state of the patient’s brain. However, the signal is easily affected by measuring instruments including electrode fault, line noise, and high electrode impedance [2], and the human body, including eye movements, blinking, heart activity, and muscle activity. [3], and then affect the quality of the signal. [4–5] (figure 1(a)[6]). The removal of physiological artifacts is more complicated than that caused by measuring instruments. It has been found in past studies that physiological artifacts may impact the collected EEG signals, greatly influencing subsequent research, as is evident in BRAIN-computer interfaces.[7] Figure 2(b)[8] shows the three most common physiological artifacts in EEG signals.

**Figure 1.**
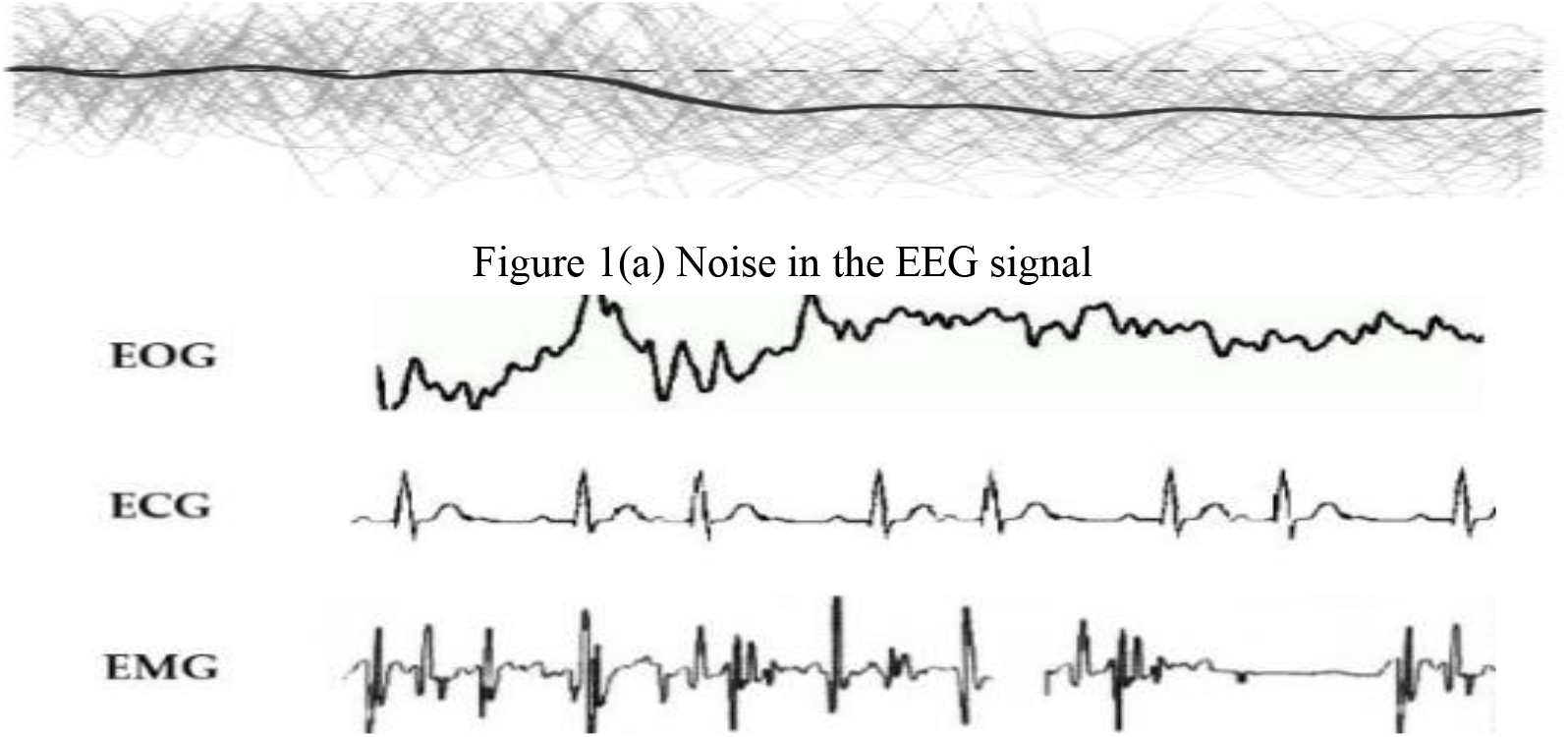
(a) Noise in the EEG signal (b) Physiological artifacts present in EEG signals.

**Figure 2.**
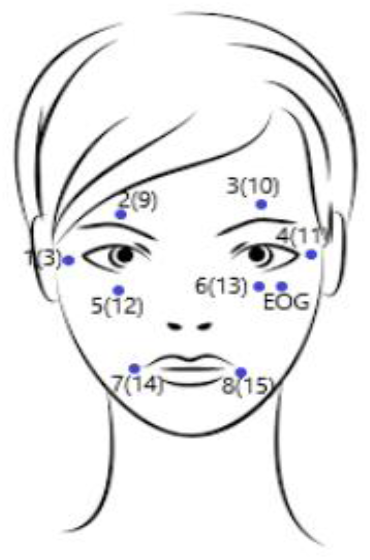
Location of external sensors

Because artifact identification and removal are the most crucial pre-processing step in clinical diagnosis or practical application, the elimination of artifacts has always been an important research content of brain neuroscience in the published papers. As we know, the Literature on artifact removal published in the past five years mainly includes algorithms based on Regression [9], Blind Source Separation (BSS) [10–12], Empirical-mode Decomposition (EMD), or Wavelet Transform algorithm to their hybrid methods [13]. However, due to the limitations of single methods such as regression and BSS, mixing methods to improve the performance of the technology Has become a potential development trend in recent years.

Whereas, up to now, there is no accurate evaluation system that can directly and accurately evaluate how well the algorithm removes artifacts from EEG signals. In previous studies, researchers mainly based on the evaluation of the EEG signal itself to indirectly assess the algorithm’s effectiveness for artifact removal [14–15]. On this basis, we propose a direct way to evaluate the effectiveness of artifacts removal methods by collecting a unique set of recorded EEG data in parallel with artifacts (EMG/ ACC/ Gyro) when EEG signals collect. Based on these external sensors, we reliably synthesized several common physiological artifacts in EEG signal acquisition. Through these physiological artifact signals, we have achieved a direct evaluation of the effectiveness of the artifact removal algorithm.

In the statistics of newly published artifact removal algorithms in the past five years, ICA [13] (independent component analysis)-related algorithms accounted for more than one-third [8]. ICA assumes that the signal source is an instantaneous linear mixture of brain sources and artificial sources, and the observed signal can be decomposed into independent components (ICs). [16]. It has been shown to work better than many other algorithms of the time and used in various studies in the field of EEG in recent years. For instance, Makeig et al. firstly applied the ICA algorithm to analyze EEG and EPR signals [17]. In 2000, Jung et al. removed artifacts from EEG by the extended ICA, and results comparing effectively to regression algorithm [18]. Romero et al. applied ICA to reduce EEG artifacts in different sleep stages and found the bidirectional property of EEG and EOG had little effect on ICA [19].

However, the decomposability of ICA to EEG signals has not been verified effectively. Whether the independent components marked as artifacts can practically represent the artifacts caused by physiological activities in the EEG signal acquisition has always been a blank. Therefore, in this project, we verified the correlation between the independent components decomposed by ICA and the artifact signals constructed through the unique data set we collected. In the end, a reliable conclusion can be drawn as to how many artifacts can ICA remove from the signal.

## Method

### ⚫ Participants

Twelve volunteers took part in the study, each receiving a varying number of electrical brain signals (3 to 8), each signal is about five minutes long. Subjects were without any known medical condition, at the beginning of participation, each subject was informed in detail about the purpose of the study and signed the written consent.

### ⚫ Experimental design

An artifact detection and removal module are indispensable for any EEG application. Even if such algorithms are intended only for offline data processing (i.e., no real-time algorithm needed), a good database of potential artifacts is needed to evaluate the respective algorithms. As far as open-source data sets, there is no reliable open-source data set that can satisfy the researchers’ research on common artifacts in EEG signals. Therefore, a direct assessment of the effectiveness of artifact removal algorithms has always been unrealistic because the current technology does not support researchers to obtain pure physiological activity signals and EEG signals directly.

In this experiment thus, we propose to create a comprehensive artifact type list firstly. After deliberations, the following set of tasks is proposed for inclusion, Resting, pressing channel, Blinking, Eye movements, Head movements, Speaking, Swallowing, clenching teeth, Frowning, and Eyebrow raise. For eye and head movements, we divide them into four different movement states, namely left and right movement, up and down movement, circular movement, and other movements. In our opinion, all the physiological activities mentioned above can contain almost all the physiological disturbances that may occur in EEG signal acquisition without introducing similar movements.

Thus, we propose to collect a set of unique data and use some external sensors to record the physiological behavior of the subject in parallel while collecting EEG signals. We reliably synthesized the artifact signals that can reflect the above-mentioned physiological activities based on these external sensors. Through these reconstructed artifact signals, we believe that they can be used to evaluate the effectiveness of the artifact removal algorithm and the decomposability of ICA to artifact components of physiological activity.

### ⚫ Experimental apparatus

Signals are collected in a bright, enclosed environment shared by both subjects and researchers. The subjects sat in comfortable chairs and performed a series of physical eye and head movements as instructed by the researchers. EEG signals were collected at a sampling rate of 512Hz, covering 31 locations [20] through Ag/ AgCI electrodes. In addition, data from 15 external sensors (Including 1 EOG sensor, 8 EMG sensors, accelerator, and gyroscope sensors) were recorded synchronously (to EEG). All subsequent artifact signals come from this unique data set. All EEG signals were referenced to electrode CPz and kept with impedance under 20kΩ. The following figures indicates the specific location of the EEG sensors. The numbers one to eight in the Figure 4 indicate the sampling position of the 8 EMG sensors. The first to sixth EMG sensors are placed around the eyes and the seventh and Eighth EMG sensors are placed around the lips.

**Figure 3.**
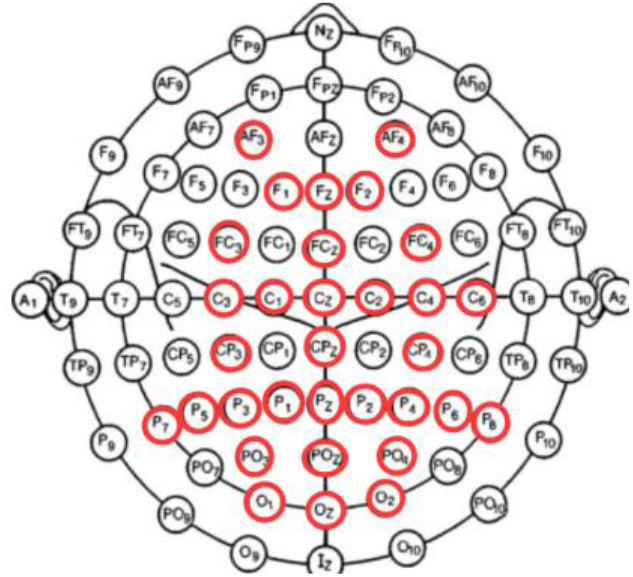
Location of EEG channel

**Figure 4.**
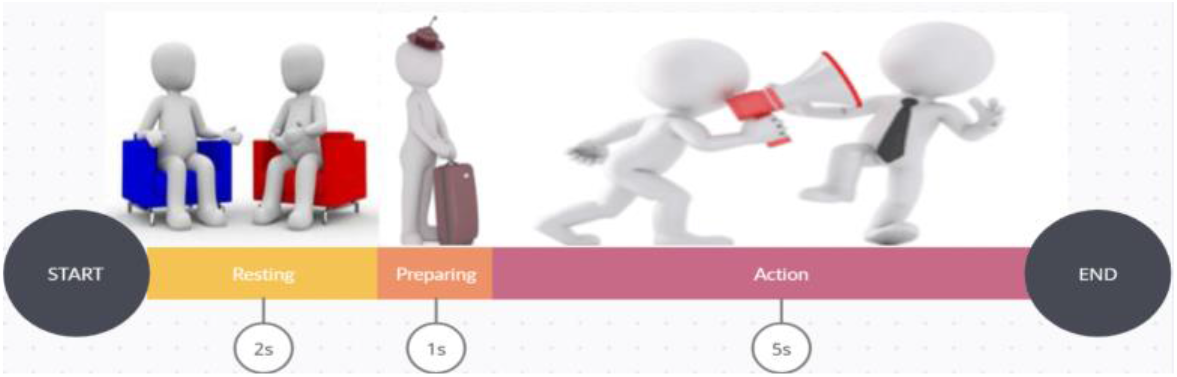
Signal collection

### ⚫ Signal collection

For the collected signals, each signal contains an independent session. There are roughly 32 repeated Trials in each session. The duration of each trial is 8 seconds consisting of three different epochs, “Resting,” “Preparing,” and “Action” period:

➢ In the “Resting” period, there are 2 seconds for the subject to rest for the upcoming test. No graphical element will be shown on the screen in this section, and subjects do not have to react just stay still.
➢ In the “Preparing” period, there is 1 second for the subject where the user is instructed to cease any artifact production and focus on the upcoming task. A cross or some other symbol on the screen will mark the beginning of this epoch.
➢ In the “Action” period, there are 5 seconds for the subject to executes the cued artifact. A text string displays on the screen the type of artifact to be executed. The disappearance of this text marks the end of the action epoch and the beginning of the “Resting” interval epoch.

All session data is collected under the above requirements, so each entire session is about 320 seconds. We collected eight such sessions for each subject to cover all the everyday physiological activities mentioned above fully. Thus, together with setup time and inter-run breaks, a complete test on a subject takes about an hour.

### ⚫ Artifact signal reconstruction

In this part, we process the signals collected by external sensors in parallel and reliably reconstruct the artifact signals corresponding to each physiological activity according to the following formulas or methods.

➢ Resting (baseline): Under ideal conditions, resting means that there is no interference in the test environment, and the signal can be approximately considered a pure signal. In this part, we asked the subjects to keep as still as possible and check whether the external sensors are triggered at the same time to obtain the nearly pure EEG signal better.
➢ Press channel: Pressing channels is one of the most common behaviors in EEG acquisition. The user (and not the operator) will randomly select a channel and press it himself. For this activity, we calculated the correlation between the 31 collected EEG signals and screened out the one with the highest correlation with the other 30 EEG signals as the artifact signal corresponding to the Press Channel.
➢ Blinking:

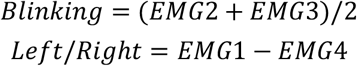
➢ Eye movements:

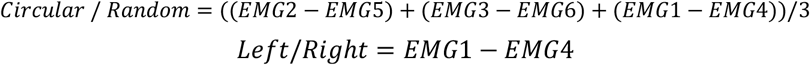
➢ Head movements:

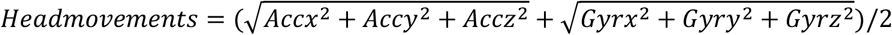
➢ Speaking:

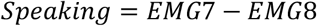
➢ Swallowing:

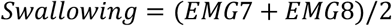
➢ Clenching teeth:

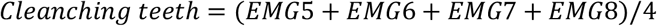
➢ Frowning:

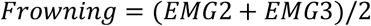
➢ Eyebrow raising:

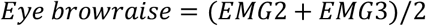

### ⚫ Benchmarked artifact removal methods selection

As a benchmark, we need to establish a systematic set of criteria to filter our benchmark methods. After, excluding naive and suboptimal PCA-based approaches and inverse approaches (too computationally expensive to apply online), we defined the following inclusion/exclusion criteria from three perspectives.

➢ Comprehensiveness: we include one method from each “category” including Regression-based, Wavelet-based, Blind-Source Separation-based (BSS) with ICA, BSS with CCA, with EMD, with SCA, Filtering methods, and Hybrid methods.
➢ Practicability: We exclude all methods that need a reference signal to work just like regression-based methods and someone only for particular artifacts, for instance, eye, muscle or movement artifacts.
➢ Availability: All of the MATLAB toolkits involved in the project are adequately free, open-source, “adequately” validated in the state-of-art and have heavily used it by far.

Therefore, on the premise of the above criteria, we selected the following artifact removal algorithms to participate in this benchmark test.

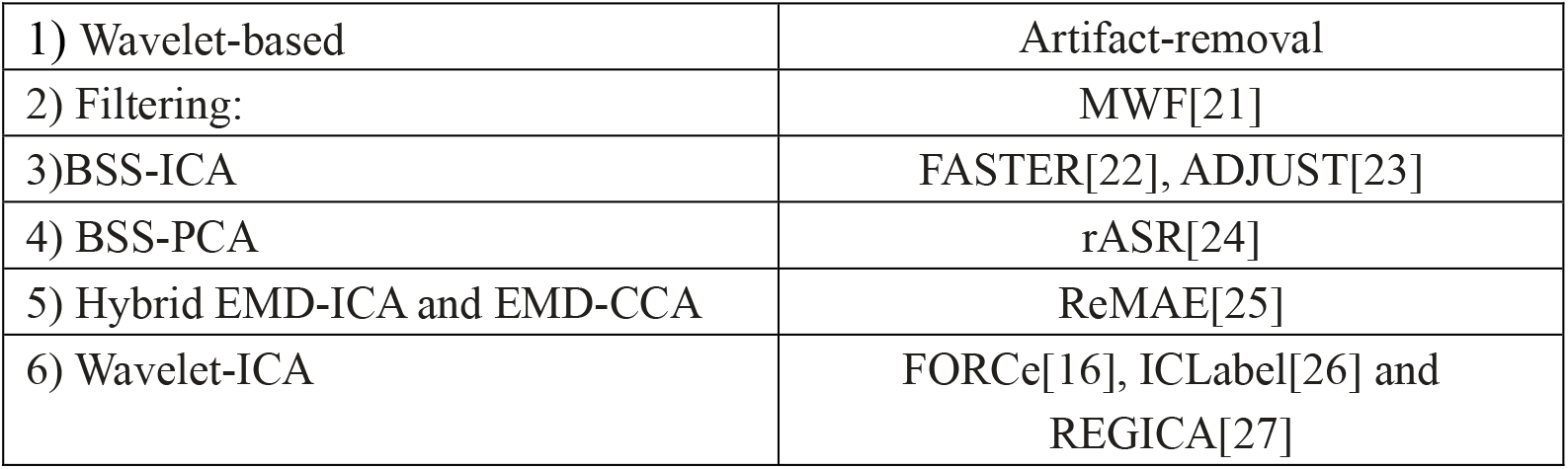

### ⚫ ICA analyzing

Independent component analysis is an algorithmic attempt to decompose a multivariate signal into independent non-Gaussian signals. Due to the multi-channel particularity of EEG signals, independent component analysis is being widely used in artifact removal algorithms. The algorithm chosen for benchmarking above reflects this very well. According to the expectations of the algorithm, independent component analysis can separate artifact signals that are affected by physiological activities during EEG signal acquisition. Does the component separated from the signal at the receiving end contain a component that can reflect the artifact signal caused by physiological activity? This is a question that needs verification. Therefore, we compare the physiological artifact signals synthesized by external sensors with the independent components separated by ICA to verify its decomposability in our design.

### ⚫ Evaluation criterion

In order to verify the ability of different algorithms to remove artifacts in EEG signals, we propose two methods: direct analysis using Pearson correlation coefficient and indirect evaluation using machine learning.

#### 1. Validation based on Pearson correlation coefficient

##### ✧ Artifact removal algorithm evaluation

The Pearson correlation [28] coefficient can directly evaluate the relationship between the synthesized artifact signal and the EEG signal. By comparing the correlation between the EEG signal before and after the artifact removal by different algorithms and the physiological artifact signal, the artifact removal ability of different algorithms is obvious.

In order to intuitively reflect the degree of influence of EEG signals by artifacts, we directly draw the correlations according to the positions of different potentials by drawing as follows, with a legend. The brighter part in the figure indicates the higher the correlation between the EEG signal at the electrodeposition and the artifact signal, and the darker the part indicates, the lower the correlation.

**Figure 5.**
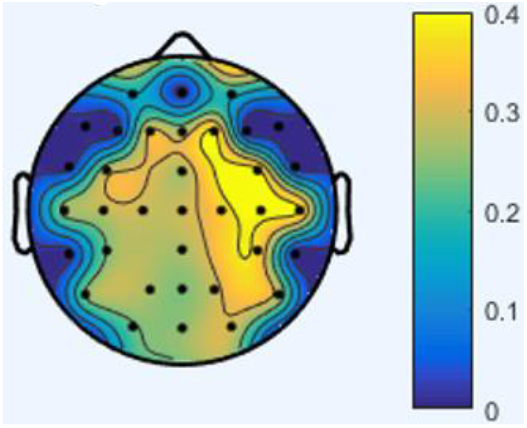
Artifact correlation

##### ✧ ICA evaluation

The verification of the ICA [29] algorithm is very similar to the evaluation of the artifact removal algorithm consider the above ten different physiological activities and based on the Pearson correlation coefficient. The difference is that in this part, correlation analysis is based on the independent components of the ICA algorithm [30] and the reconstruction of artifact signals for each physiological activity. This correlation can be regarded as the extent to which the separated independent components can restore the reconstructed artifact signal.

Since independent component analysis can restore various components other than the EEG signal based on the multi-channel EEG signal, we only save the independent component with the highest correlation with the reconstructed artifact signal as the artifact component. At the same time, we use the artifact evaluation mechanism of the algorithm with the best artifact removal effect above to remove the artifact components and recalculate the correlation between the independent components that do not contain the artifact components and the reconstructed artifact signal.

By analyzing the correlation before and after removing the artifact components, specific judgments can be made on the effect of ICA in removing the artifact components. In addition, we also classified each set of related data according to different physiological activities and further subdivided the ICA’s ability to separate artifacts through standard deviation [31] (Z-Score) for each physiological behavior.

#### 2. Artifact Classification Based on Machine Learning

As a benchmark, we tried to cite machine learning [32] as an additional corroboration method. In order to verify whether the physiological artifacts in the signal can be effectively removed, we try to classify EEG signals in 9 different physiological states (without Resting). Under ideal conditions, if the algorithm effectively removes the artifacts in the signal, the model should not be able to present a blind classification of the cleared signal, so that the classification accuracy [33] should be much lower than that of the uncleared signal. Therefore, we built a machine learning model according to the following steps. In order to control the experimental variables only for the signals cleaned by different algorithms, all models and parameters in this part are precisely the same, and all algorithms use the same training/testing set.

##### ✧ Data processing: Removing baseline noise by wavelet transform

When collecting EEG signals, baseline drift, as a low-frequency artifact, may be caused by poor electrode contact that affects electrode impedance. [34] This project uses the characteristics of multi-scale and multi-resolution of wavelet transform to select appropriate wavelet transform [35] function and decomposition level to decompose the signal in multi-scale. Since the main component of baseline drift is a gradually changing trend component, it will directly appear on a larger scale in the wavelet decomposition. As long as the component at this scale is directly removed during the reconstruction process, the baseline correction can be achieved.

##### ✧ Feature extraction: Extract the proportion of wavelet energy

The essence of the wavelet transform is the filtering process of the original signal, and the decomposition results are different with the different selection of the wavelet function [36]. However, no matter how the wavelet function selects the centre frequency and bandwidth of the filter used for each decomposition into a fixed proportion, that is, it has the so-called “constant Q” characteristic. Therefore, the smoothing signals and detail signals in each scale space can provide the local information of the original signal in time domain, especially the signal composition information in different frequency bands. If the energy of signals in different decomposed frequency bands is solved, these energy values can be arranged in order of frequency bands to form characteristic energy for identification.

In this project, firstly, the EEG signals are stratified by wavelet packet decomposition, and then the binary wavelet transform is used to extract the energy distribution features on the scale and space. Wherein, the wavelet energy of frequency band *i* of layer *j* can be calculated by using wavelet transform as follows:

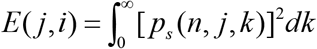

In the formula, *p*_*s*_ (*n*, *j*, *k*) is the wavelet packet transform coefficient.

The total energy of bone vibration signal *E*_sum_ can be obtained by summing the wavelet energy of each frequency band:

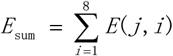

The percentage of the wavelet energy of each frequency band in the total energy is called the wavelet energy ratio:

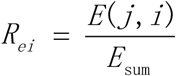

In addition, we also extracted kurtosis, skewness, variance, standard deviation and other common features for EEG signal classification.

##### ✧ Artifact classification: XGBoost[37]

As an excellent machine learning algorithm, XGBoost is widely used in various machine learning classification problems, with highly efficient, flexible, and portable characteristics. After comparing the running results and the time spent by different algorithms, we finally choose XGBoost for signal classification verification. In order to more intuitively reflect the classification results [38], we not only compared the classification accuracy of the algorithms but also plotted their confusion matrix [39]. By comparing the classification results of the signals after cleaning by different algorithms and the classification results of the original uncleaned signals, we can intuitively compare different algorithms For the effectiveness of artifact removal.

### ⚫ Data use

No participants were deleted in the analysis reported in this manuscript. All results are based on either the saved original EEG data or the processed feature vector. Due to technical problems, some signal lengths are much smaller than the rated signal length and are therefore regarded as error signals and removed from the data set.

## Result

### ⚫ Analysis of the influence of artifacts on uncleaned signals

In order to set up the control group, we analyzed the original EEG signals, classified different physiological activities, and calculated the correlation of the artifact signals based on the physiological activity. The following results are average correlations for each physiological behavior based on all samples. The characteristics it reflects are universally persuasive.

The brighter the image, the greater the influence of the physiological activity on the EEG signal sampling, and the darker the image, the less the influence of the EEG signal sampling. It is important to note that the degree of influence of these physiological behaviors can be referred to the colormap on the right for specific EEG electrodes.

**Figure 6.**
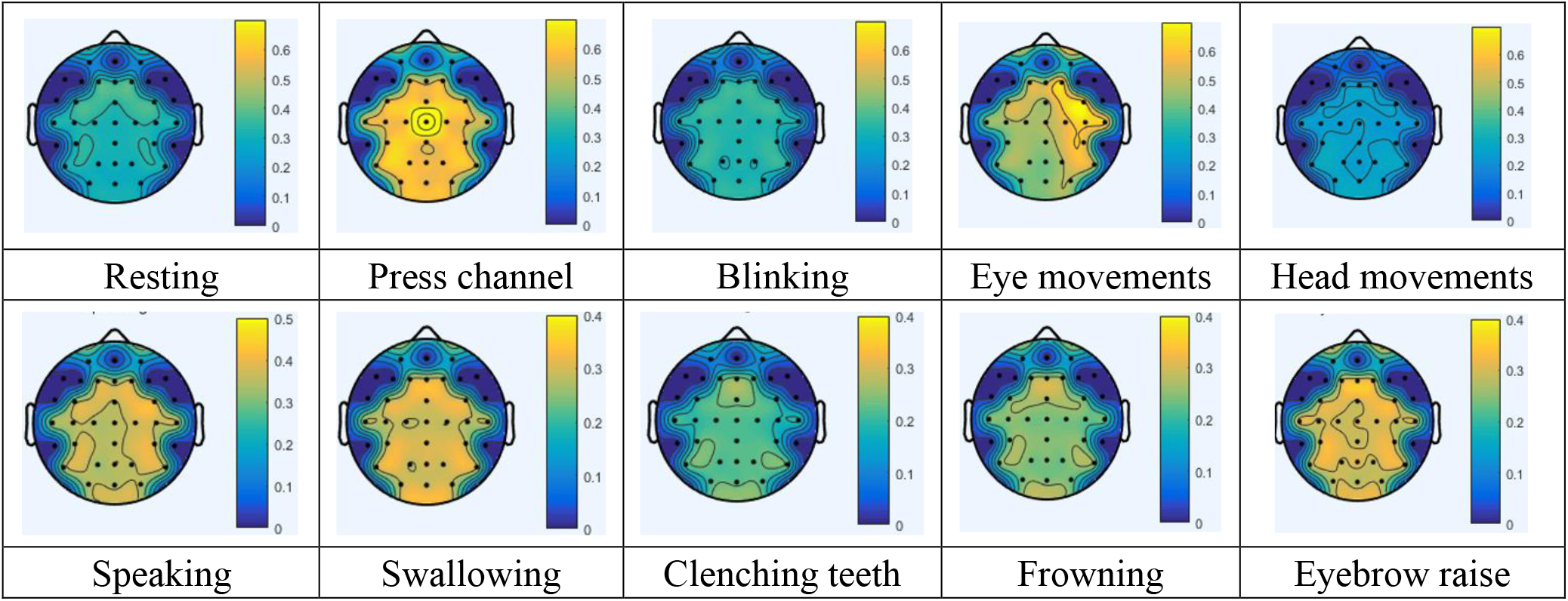
The influence degree of different physiological activities on EEG signal

These artifact correlations conform to our predictions, and it is not difficult to find that the correlation of “Resting” is almost less than 0.3 as for the non-artifact “Resting” period set by us. The correlation coefficient of local “Eye movement” is the highest among all physical activities (much higher than 0.6), while the correlation of most other activities is around 0.4. As for “Blinking” and “Head movement,” their correlation is close to “Resting,” which is somewhat unexpected and may be related to the calculation method. The influence of “Eyebrow raising,” “Speaking,” and “Swallowing” on the collected EEG signals is also severe; the degree of influence is very similar for different electrode positions, ranging from 0.3 to 0.4. Thus, on the whole, different activities and “Resting” nearly present a contrasting situation.

### ⚫ Overview of cleaning effectiveness of different algorithms

In order to intuitively compare the ability of different algorithms to remove artifacts, we have unified the original EEG signals and the EEG signals after the artifacts have been cleaned by different algorithms and have summarized the EEG signals containing different physiological activities and generated a total of 11 images. Overview picture (one original signal and ten effect pictures of different artifact removal algorithms).

**Figure 7.**
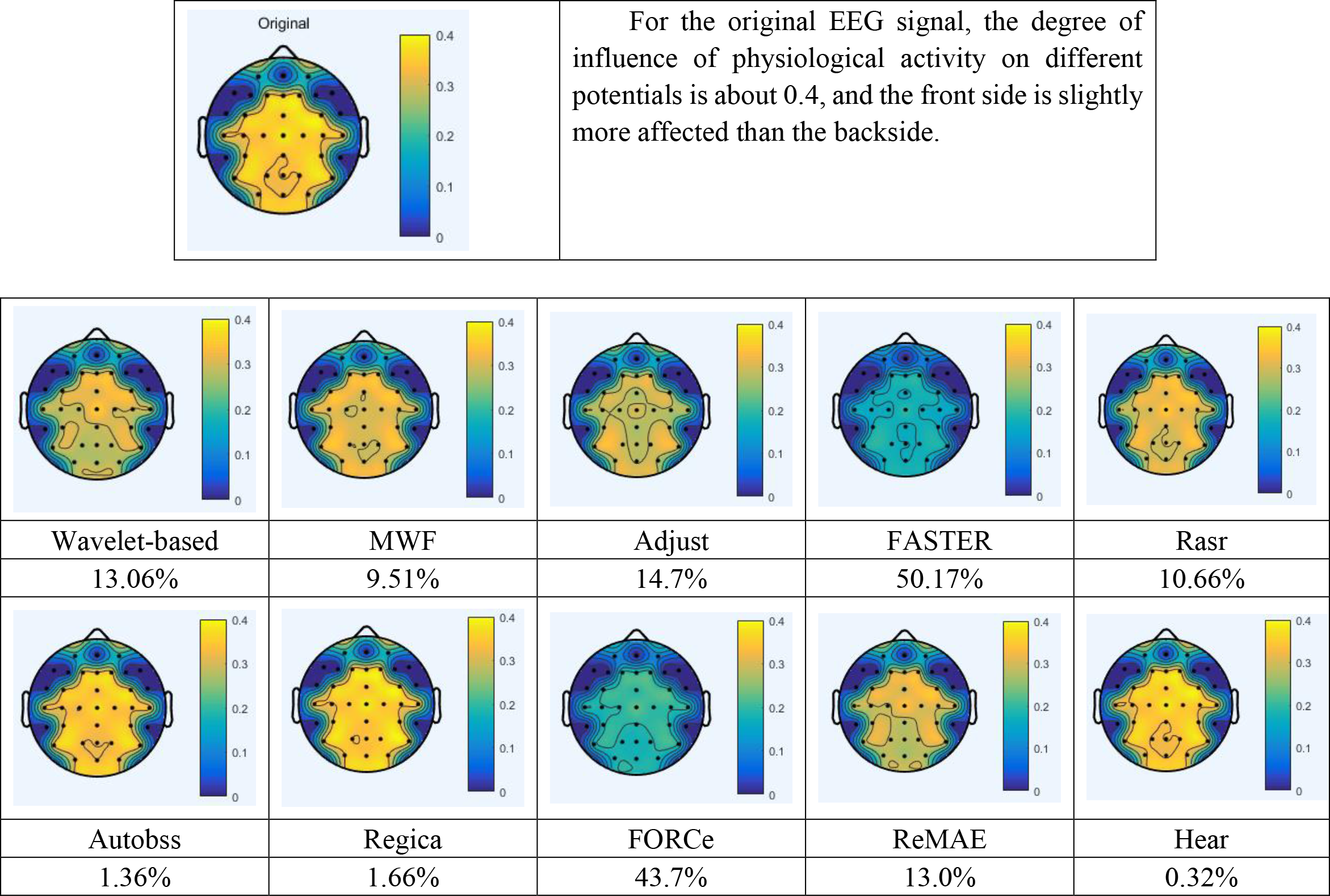
Comparison of artifact removal effects of different algorithms

The table shows the artifact removal effect diagrams of different algorithms and the removal ratio relative to the artifact correlation of the original signal explicitly. The artifact removal effects of FASTER and FORCe are relatively excellent (50.17% and 43.7%, respectively). Artifacts significantly reduce the EEG signal after cleaning, and the correlation of almost every potential is lower than 0.2. The artifact removal effect of Wavelet-based, MWF, Rasr, and ReMAE is relatively close to about 10%. Compared with the uncleaned EEG signal, although the correlation is significantly lower. There is still a noticeable difference compared with FASTER and FORCe. For Autobss, Regica, and Hear, the artifact removal effect of the overview is not very obvious. The signal correlation after removing the artifacts is closer to the original signal, mainly because the artifact removal of these algorithms is more targeted. Some of the physiological behaviors evaluated this time are not within the range of artifacts removed.

### ⚫ The difference after cleaning

In order to observe the artifact removal effect of the artifact removal algorithm on different physiological behaviors, we divided the data according to different physiological activities. The following figure shows the analysis of the artifact removal effects of FORCe and FASTER, which have the best artifact removal effect for ten different physiological behaviors. The brighter the image, the better the algorithm can remove artifacts from the current electrode position. The specific degree of removal can refer to the colormap on the right

**Figure 8.**
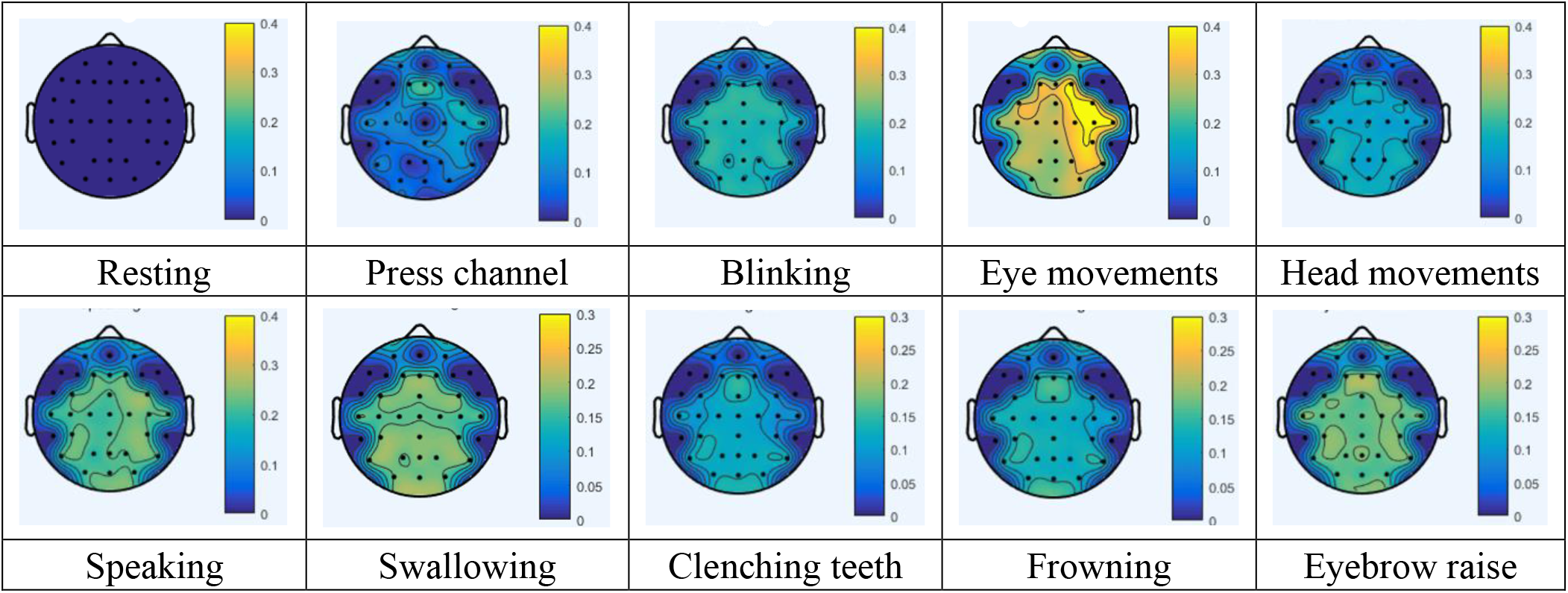
The artifact removal ability of FORCe

**Figure 9.**
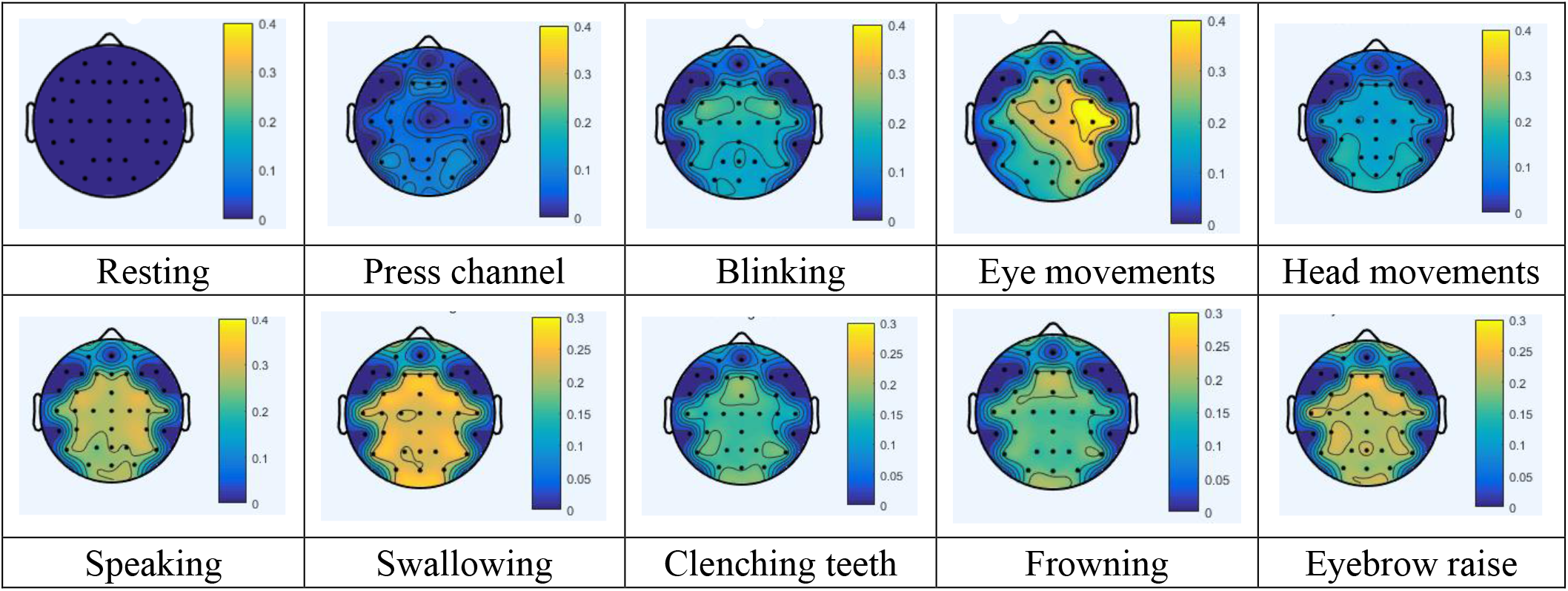
The artifact removal ability of FASTER

In general, for different physiological activities, these two artifact removal algorithms have apparent effects. For “Speaking,” “Swallowing,” “Clenching teeth,” “Frowning,” “Eyebrow raise” these physiological activities, FASTER’s artifact removal effect has a slight lead. However, although FASTER outperforms FORCe in the overview in the table above, FORCe performs even better for “Press channel,” “Blinking,” and “Eye movement” activities. The two algorithms have almost the same degree of cleaning for “Head movement”. “Resting” is always regarded as static without artifacts, so no analysis and statistics are made in this part.

**Figure 10.**
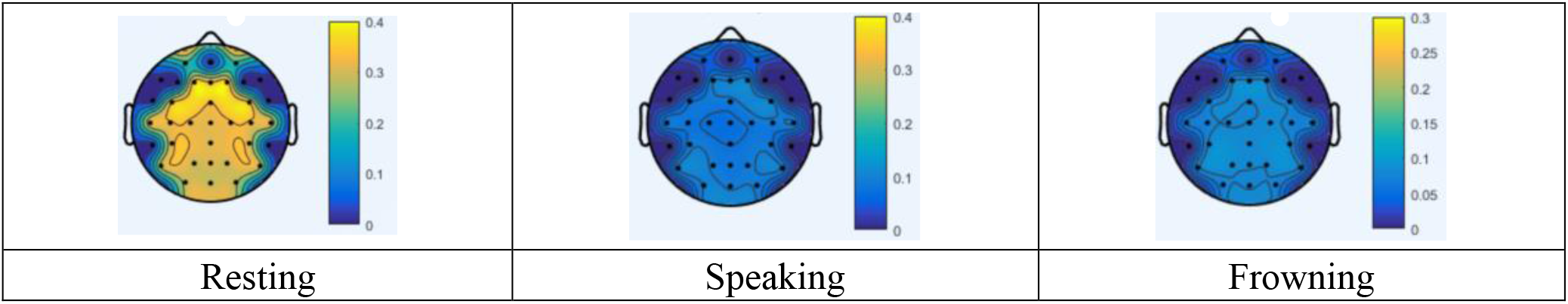
Partial physiological activity artifact correlation after FASTER processing

It is not difficult to see from the figure that in many physiological activities of the processed EEG signals, the correlation between the corresponding artifact signals and EEG signals at different electrode positions was even much lower than that in the untreated “Resting” state. Therefore, it is not difficult to see that the artifact removal effect is pronounced.

### ⚫ The machine learning classification results

It is not difficult to see from the classification results that the cleaned signals made models have difficulty classifying the subjects. This is well reflected in the confusion matrix shown below for uncleaned signals, and FORCe cleaned signals. The classification results were mainly distributed on the diagonal for unwashed EEG signals, whereas for FORCe washed signals, this phenomenon was much weaker, more like a random distribution. The accuracy of other model classification results is also shown in detail in the following table.

**Figure 11.**
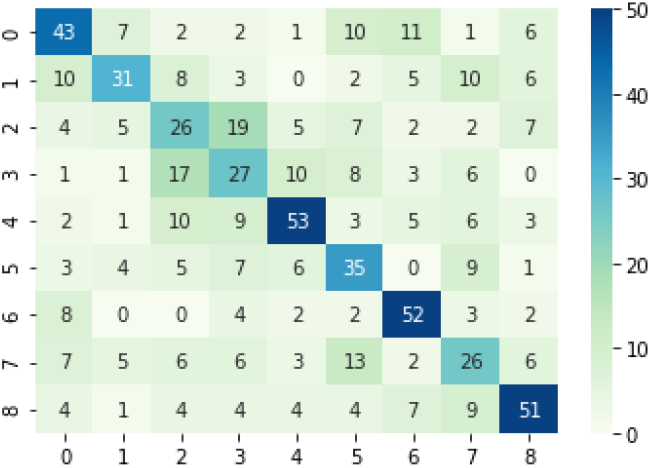
Classification results of unwashed EEG signals

**Figure 12.**
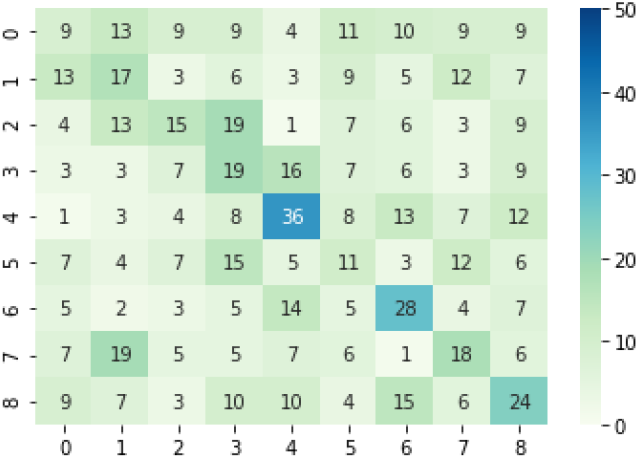
Classification results of FORCe cleaned EEG signals

**Table 1.** classification accuracy for different method

| Method | Result |
| --- | --- |
| Uncleaned | 0.4879 |
| Wavelet-based | 0.4355 |
| MWF | 0.3758 |
| Adjust | 0.3390 |
| FASTER | 0.4709 |
| Rasr | 0.4567 |
| Autobss | 0.3872 |
| FORCE | 0.2510 |
| ReMAE | 0.3220 |
| Hear | 0.4624 |

Although the algorithms other than FORCe also reduced the classification accuracy to varying degrees, there was still a particular gap with FORCe. MWF, Adjust, Autobss, and ReMAE have more obvious artifact removal effects than other algorithms. Surprisingly, The classification accuracy of FASTER was very close to that of the unwashed signal, contrary to the excellent results obtained by correlation analysis in the last part.

### ⚫ Verify independent component analysis

#### ➢ Based on correlation

To verify the ICA algorithm, we performed independent component analysis on the EEG signals containing artifacts, which, in theory, could separate the artifacts from the EEG signals. Based on the analysis results of the first two parts, although both FORCe and FASTER contain ICA algorithm, we finally referred to the discrimination method of independent components in FORCe and compared the highest correlation between artifact signals and all components before and after removing this component. After analyzing the independent components of ICA, considering all physiological behaviors, we found that the highest independent component correlation with artifact signal was more than 0.7. After removing the components judged to be artifacts using the artifact elimination algorithm in FORCe, the correlation is reduced to 0.3. For each physiological behavior, we also made independent calculations. The specific results are shown in the table below, which are very close to the average results.

**Table 2.**
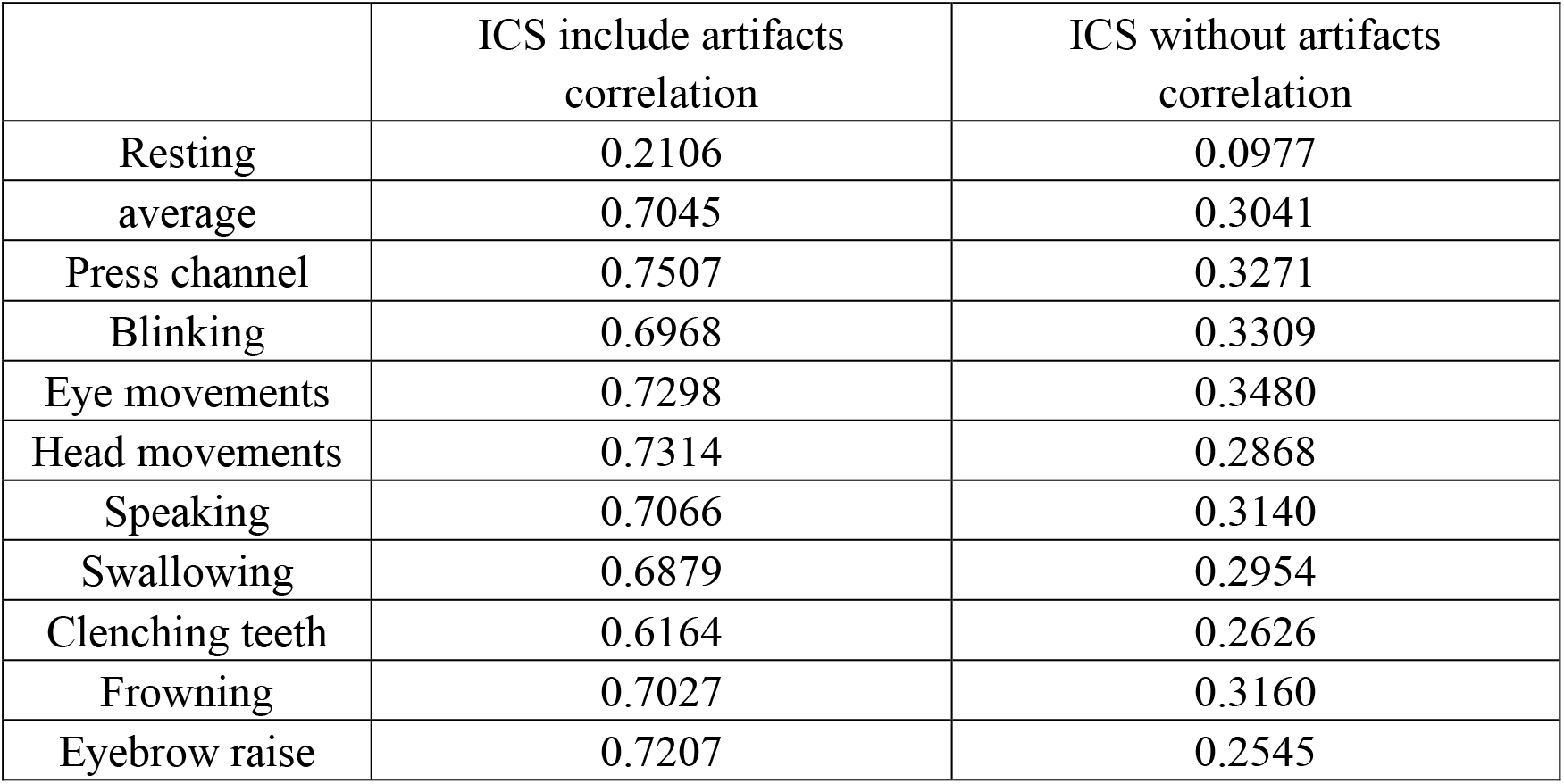
The separation degree of artifacts of different physiological activities by ICA(①)

In the above table, Resting is regarded as the EEG signal without artifact in the ideal state, and before cleaning, its correlation (0.21) is far lower than that of other physiological states. After eliminating artifact components, the correlation is reduced to 0.09.

**Table 3.** The separation degree of artifacts of different physiological activities by ICA(②)

|  | ICS include artifacts<br>correlation | ICS without artifacts<br>correlation |
| --- | --- | --- |
| average | 0.7787 | 0.2031 |
| Press channel | 0.944 | 0.357 |
| Blinking | 0.8543 | 0.1287 |
| Eye movements | 0.984 | 0.2328 |
| Head movements | 0.7889 | 0.2826 |
| Speaking | 0.662 | 0.1447 |
| Swallowing | 0.7173 | 0.3193 |
| Clenching teeth | 0.586 | 0.1318 |
| Frowning | 0.7413 | 0.1149 |
| Eyebrow raise | 0.7278 | 0.1119 |

What is most noteworthy is that in all the sample data, the data of some subjects are analyzed, and the results are excellent. The artifact components of many physiological behaviors are even close to 1, and the complete results are also close to 0.8. After removing the artifact component, the component with the highest correlation between the remaining component and the artifact signal mainly maintains at 0.1. After removing the artifact component, the correlation means of all physiological activities also decreased significantly. This result’s physiological activity is very close to what we would assume under ideal conditions.

#### ➢ Base on Z-Score

Based on the results in Table 4, we further standardized the data by using the Z-score, the independent component of Resting after eliminating fake artifacts is taken as the benchmark, and the specific results are shown in Table 5. The values in the table can be viewed as the number of unit standard deviations from the benchmark.

**Table 4.**
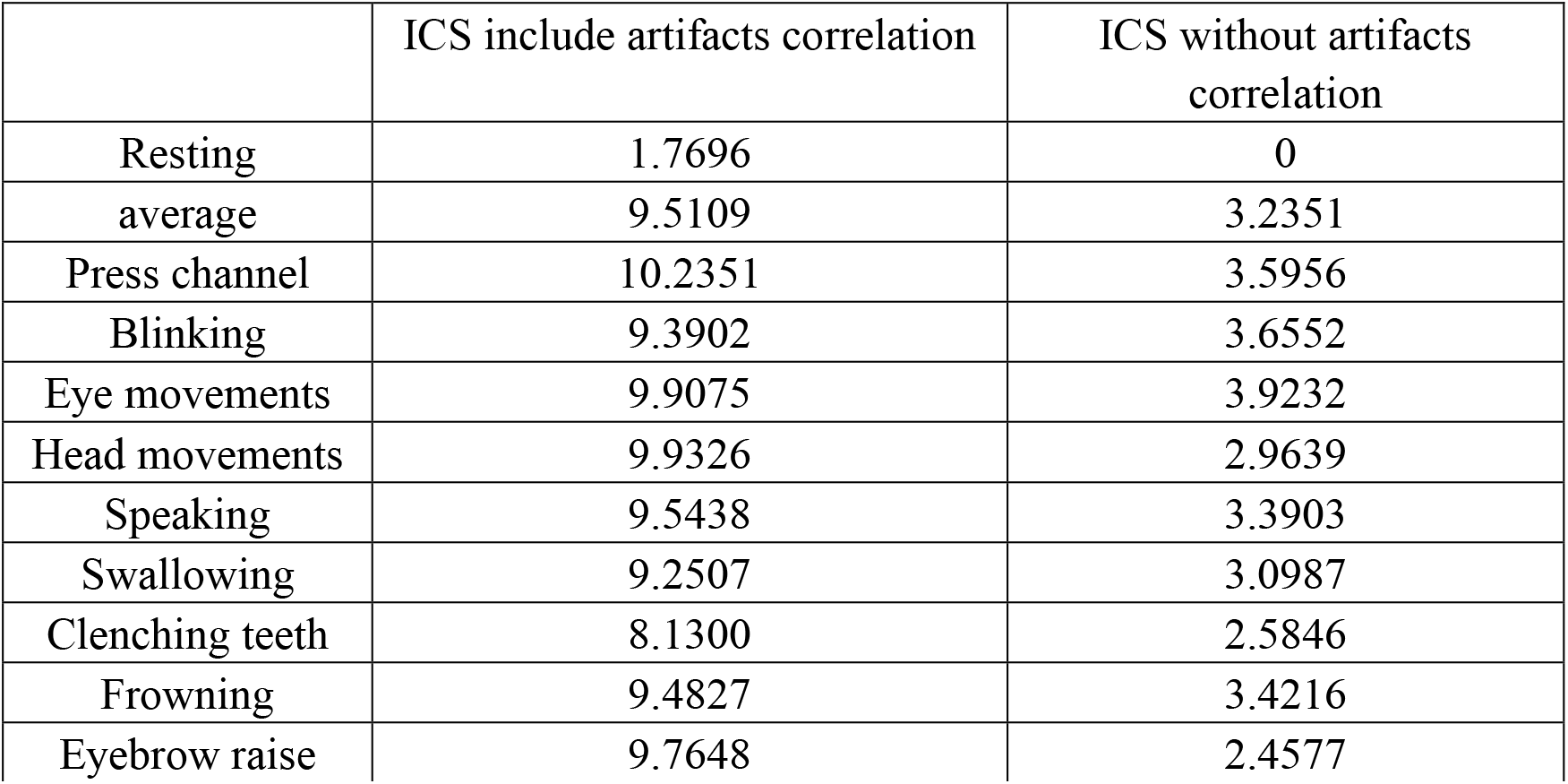
ICA Z-score standardization results

As can be seen from the table, under the same Resting state, the calculated result including artifact components is “closest” to the benchmark state, with only a difference of less than two standard deviations. In addition, the Z-score of different physiological activities with artifact components removed was far less than that without artifact components removed.

## Conclusions

In this benchmark test, we analyzed several popular artifact removal algorithms. The correlation results of different algorithms are different. Based on the independent component analysis, FORCe and FASTER have the best effect and excellent artifact removal effect for each physiological activity, especially the most relevant “Eye movement.” Almost 40% of the artifacts are removed. For other physiological activities, no less than 20% of artifacts were removed. For other removal methods, Hear, Autobss, and regica mainly only focus on a particular activity, especially eye movements. Furthermore, the removal effect of rASR, MWF, and others, after the artifact correlation was removed by 10%-20%, pronounced the cleaning effect on the general diagram. Overall, these algorithms are much better at removing the subtle physiological movements of the face than the artifacts generated by head movements. In general, compared with facial artifacts, eye artifacts are easier to remove.

In the analysis based on machine learning, the classification results presented by different algorithms are generally very similar to our expectations. After cleaning, the EEG signals bring varying degrees of difficulty to model classification, resulting in a decrease in classification accuracy compared with the uncleaned signals. FORCe still showed excellent performance, with classification accuracy decreasing by about 40%, similar to the previous analysis based on correlation. However, the result of FASTER is not as good as in the previous section. It may be because the effect of FASTER in removing artifacts caused by environmental factors is better than that caused by the subjects’ physiological behaviors.

Due to FORCe ‘s excellent performance in benchmark tests, we adopted his artifact component discrimination mechanism. In the original hypothesis, we initially believed that the independent component containing artifacts had the highest correlation of more than 0.9. Under ideal conditions, the correlation should be close to 0 after eliminating these components containing artifacts. While Resting, as a physiological state without artifacts, the correlation between its independent component and artifact signal should also be close to 0. However, after analysis, it is not difficult to find that the overall result reflects our expected results objectively; the results based on partial sample data reflect this point well Even in some physiological activities, the results are not the same as what we assume under ideal conditions. Compared to the expected complete elimination of all artifact components, ICA eliminated almost 70% of artifacts. Although it still has a gap with the ideal state of completely stripping the artifacts, it is still an excellent EEG artifact removal technology.

By verifying the results of the Z-score, we find that ICA can make the original signal closer to the “Resting” state, which does not contain artifacts at all under the ideal state, by eliminating the components regarded as artifacts. Furthermore, after analyzing the standard deviation after Z-score standardization, for each physiological state, its artifact elimination effect is similar to the previous result based on correlation, which can eliminate more than 70% artifacts in the signal.

